# Increased posterior default mode network activity and structural connectivity in young adult *APOE*-ε4 carriers: a multi-modal imaging investigation

**DOI:** 10.1101/285536

**Authors:** Carl J. Hodgetts, Jonathan P. Shine, Huw Williams, Mark Postans, Rebecca Sims, Julie Williams, Andrew D. Lawrence, Kim S. Graham

## Abstract

Young adult *APOE*-ε4 carriers show increased activity in posterior regions of the default mode network (pDMN), but how this is related to structural connectivity is unknown. Thirty young adults (half *APOE*-ε4 carriers, the other half *APOE*-ε3ε3/ε2ε3; mean age 20 years) were scanned using both diffusion and functional magnetic resonance imaging. Diffusion tractography was used to quantify the microstructure (mean diffusivity, MD; fractional anisotropy, FA) of the parahippocampal cingulum bundle (PHCB), which links pDMN and the medial temporal lobe. *APOE*-ε4 carriers had lower MD and higher FA relative to non-carriers in PHCB. Further, PHCB microstructure was selectively associated with pDMN activity during a scene discrimination task known to be sensitive to Alzheimer’s disease (AD). These findings are consistent with a lifespan view of AD risk, where early-life structural and functional brain changes in specific, vulnerable networks leads to increased neural activity that may ultimately trigger amyloid-ß deposition.

## 1. Background

The default mode network (DMN) is a large-scale brain system displaying continuously high levels of coordinated activity in the resting-state (Raichle, 2015). DMN activity is maintained or enhanced during internally directed cognition (e.g., autobiographical memory, mind-wandering), and is attenuated during many cognitive tasks demanding external perceptual attention (e.g. episodic memory encoding) (Raichle, 2015).

Rather than constituting a single, unitary brain network, the DMN can be divided into several functionally dissociable subsystems (Andrews-Hanna et al., 2010; Raichle, 2015), which are affected differently by Alzheimer’s disease (AD) progression (Myers et al., 2014). Notably, the posterior DMN (pDMN), comprising posterior cingulate, precuneus and retrosplenial cortex (Cauda et al., 2010), is one of the earliest brain areas to undergo amyloid-ß accumulation and reduced metabolism in AD (Gonneaud et al., 2016; Palmqvist et al., 2017). The pDMN also constitutes the brain’s structural “core” and is characterized both by high levels of baseline activity/metabolism and dense functional and structural inter-connectivity (Bero et al., 2012; Buckner et al., 2009; Hagmann et al., 2008).

The striking spatial overlap between the pDMN and regions that show early amyloid-ß accumulation has led to a ‘lifespan systems vulnerability’ (LSV) account of AD, where increased activity and connectivity - over the *lifetime* - may predispose this region to later-life amyloid-ß (Buckner et al., 2009; de Haan et al., 2012; Jagust and Mormino, 2012). Providing strong evidence for a link between neural activity/connectivity and amyloid-ß, a study in transgenic mice (Bero et al., 2011) reported that interstitial amyloid-ß levels were associated with increased markers of neural activity, and this, in turn, predicted amyloid-ß deposition, particularly in pDMN ((Bero et al., 2011); see also (Yamamoto et al., 2015)). A further study showed that region-specific levels of functional connectivity in young Aß-mice was proportional to the degree of amyloid-ß burden in older animals (Bero et al., 2012). Similarly, human neuroimaging studies have reported associations, within-subjects, between the degree of baseline pDMN functional connectivity and subsequent amyloid-ß load in both mild cognitive impairment (MCI) (Myers et al., 2014) and cognitively-normal older adults (Jack and Holtzman, 2013). While these studies suggest that functional properties may drive pathological markers, it could reflect a later-life compensatory response induced by early amyloid-ß burden in key networks (Jagust and Mormino, 2012; Jones et al., 2016).

If later-life amyloid-ß deposition in pDMN is associated with increased functional activity/connectivity across the lifespan, then alterations may be evident in younger individuals at elevated risk of AD. Further, as amyloid-ß deposition is highly unlikely in young adults (de Haan et al., 2012; Mormino, 2014), this approach addresses a key limitation of studies in elderly individuals where compensatory functional activity may reflect early pathology (Jagust and Mormino, 2012).

The *APOE*-ε4 allelle is the strongest genetic risk factor for both sporadic early and late-onset AD (Liu et al., 2013) and is strongly linked to amyloid-ß accumulation in later life (Gonneaud et al., 2016). Functional magnetic resonance imaging (fMRI) studies have typically found that young *APOE*-ε4 carriers show increased activity, relative to non-carriers, in pDMN and inter-connected medial temporal lobe (MTL) regions (Dennis et al., 2010; Filippini et al., 2009; Shine et al., 2015). *APOE*-ε4 carriers have also been shown to have greater intrinsic functional connectivity in the DMN during “rest” (Filippini et al., 2009).

Given the view that pDMN vulnerability to amyloid-ß accumulation is linked to its role as a large-scale connectivity hub (Jagust and Mormino, 2012; Jones et al., 2016), the heightened pDMN activity in young *APOE*-ε4 carriers may itself be linked to variation in structural connectivity (Brown et al., 2011; de Haan et al., 2012) - particularly those connections linking pDMN with other regions affected early in AD.

The major white matter connection linking pDMN with MTL (particularly parahippocampal cortex) (Greicius et al., 2009; Heilbronner and Haber, 2014) is the parahippocampal cingulum bundle (PHCB). Studies applying diffusion magnetic resonance imaging (dMRI) – a method allowing *in vivo* quantification of white matter microstructure – have reported greater mean diffusivity (MD) and lower fractional anisotropy (FA) in the PHCB of cognitively-normal older *APOE*-ε4 carriers compared to non-carriers (Heise et al., 2014). One cross-sectional dMRI study found that young adult *APOE*-ε4 carriers had higher PHCB FA but showed a steeper decline across life, leading to relative FA reduction relative to non-carriers from mid-life onwards ((Felsky, 2013); see also (Brown et al., 2011)). Disruption of this pathway is also seen in both MCI and AD (Mito et al., 2018; Rieckmann et al., 2016), and has been linked to pDMN activity/metabolism in AD (Villain et al., 2008) and amyloid-ß burden in preclinical AD (Racine et al., 2014). Overall, these studies point toward the potential early-life vulnerability of a broader posterior network that is structurally underpinned by the PHCB (Greicius et al., 2009). Further, they support the view that increased brain activity over the lifespan – driven by structural connectivity - leads to amyloid-ß deposition, and ultimately results in decreased activity/connectivity in later life due to “wear and tear” (Jagust and Mormino, 2012).

It is unclear, however, whether these PHCB microstructural alterations are evident earlier in life when amyloid-ß deposition is unlikely, concomitant with the identified functional changes in college-aged adults (Dennis et al., 2010; Filippini et al., 2009; Shine et al., 2015). Moreover, if increased activity in pDMN stems from its role as a large-scale connectivity hub (Brown et al., 2011; de Haan et al., 2012), then those individuals who show elevated pDMN activity (Filippini et al., 2009; Shine et al., 2015) should also have “increased” structural connectivity (de Haan et al., 2012). To address these questions we applied high angular resolution dMRI (HARDI (Tuch et al., 2002)), alongside constrained spherical deconvolution (CSD) tractography (Jeurissen et al., 2011), to test whether the presence of an *APOE*-ε4 allele in young adults, who are unlikely to harbor amyloid burden, influences PHCB tissue microstructure. Given evidence that young *APOE*-ε4 carriers show elevated pDMN activity at rest and during tasks (Shine et al., 2015), we predicted that *APOE*-ε4 carriers would show greater FA and lower MD in the PHCB, compared to non-carriers. Finally, to demonstrate a link between activity and connectivity, as predicted by an LSV view of AD risk, we examined whether inter-individual variation in PHCB tissue microstructure was associated with pDMN activity during a scene discrimination task that is sensitive to early AD (Lee, 2006).

## 2. Material and Methods

### 2.1. Participants

Full details regarding DNA extraction and genotypic distribution can be found in a previous article (Shine et al., 2015). The available sample for this analysis was 30 participants (15 per group; 14 females per group) - a sample size similar to other structural/functional studies of *APOE*-ε4 (Dennis et al., 2010; Filippini et al., 2009; Oh and Jagust, 2013). The non-carrier *APOE* allele distribution was 10 *APOE*-ε3ε3 and 5 *APOE*-ε2ε3 individuals. The carrier *APOE* allele distribution was 14 *APOE*-ε3ε4 and 1 *APOE*-ε2ε4. Both groups were matched for age (carriers: 19.7 years, S.D. = 0.84; non-carriers: 19.7 years, S.D. = 0.89) and education level. Family history was matched across the groups, with two reports of a positive family history in each. All participants were right-handed, native English speakers with normal or corrected-to-normal vision, and had no self-reported history of neurological/psychiatric disorders. Groups were well matched on standard neuropsychological tests (Shine et al., 2015). All experimental procedures were conducted in accordance with, and were approved by, the Cardiff University School of Psychology Research Ethics Committee. Informed consent was obtained from all participants, and research was conducted in a double-blind manner.

### 2.2. MRI scan parameters

Imaging data were collected at the Cardiff University Brain Research Imaging Centre (CUBRIC) using a GE 3-T HDx MRI system (General Electric Healthcare, Milwaukee, WI) with an 8-channel receive-only head coil. Whole brain HARDI (Tuch et al., 2002) data were acquired using a diffusion-weighted single-shot spin-echo echo-planar imaging (EPI) pulse sequence with the following parameters: TE = 87ms; voxel dimensions = 2.4 × 2.4 × 2.4 mm^3^; field of view = 23 × 23 cm^2^; 96 × 96 acquisition matrix; 60 slices (oblique-axial with 2.4 mm thickness). Acquisitions were cardiac gated using a peripheral pulse oximeter. Gradients were applied along 30 isotropic directions with b = 1200 s/mm^2^. Three non-diffusion weighted images were acquired with b = 0 s/mm^2^.

### 2.3. Diffusion MRI preprocessing

Motion and eddy current correction was conducted using ExploreDTI (Leemans and Jones, 2009). Partial volume corrected maps of tissue FA and MD were generated by applying the bi-tensor ‘Free Water Elimination’ (FWE) procedure (Pasternak et al., 2009). FA reflects the extent to which diffusion is anisotropic, or constrained along a single axis, and can range from 0 (fully isotropic) to 1 (fully anisotropic). MD (10^−3^mm^2^s^-1^) reflects a combined average of axial (diffusion along the principal axis) and radial diffusion (diffusion along the orthogonal direction).

### 2.4. Tractography

Deterministic whole-brain tractography was conducted in ExploreDTI (Leemans and Jones, 2009) using the CSD model (Jeurissen et al., 2011), which extracts multiple peaks in the fiber orientation density function (fODF) (Vettel et al., 2017). Streamlines were reconstructed using the following parameters: fODF amplitude threshold = 0.1; step size = 0.5 mm; angle threshold = 60°).

Three-dimensional reconstructions of the PHCB (Figure 1A) were obtained from individual subjects using a Boolean, way-point region of interest (ROI) approach, where “AND” and “NOT” ROIs were applied and combined to isolate PHCB streamlines in each subject’s whole-brain tractography data. These ROIs were drawn manually on the direction-encoded FA maps in native space by one experimenter (HW) who was blind to *APOE*-ε4 carrier status and quality-assessed by a second experimenter (CJH).

**Figure 1.**
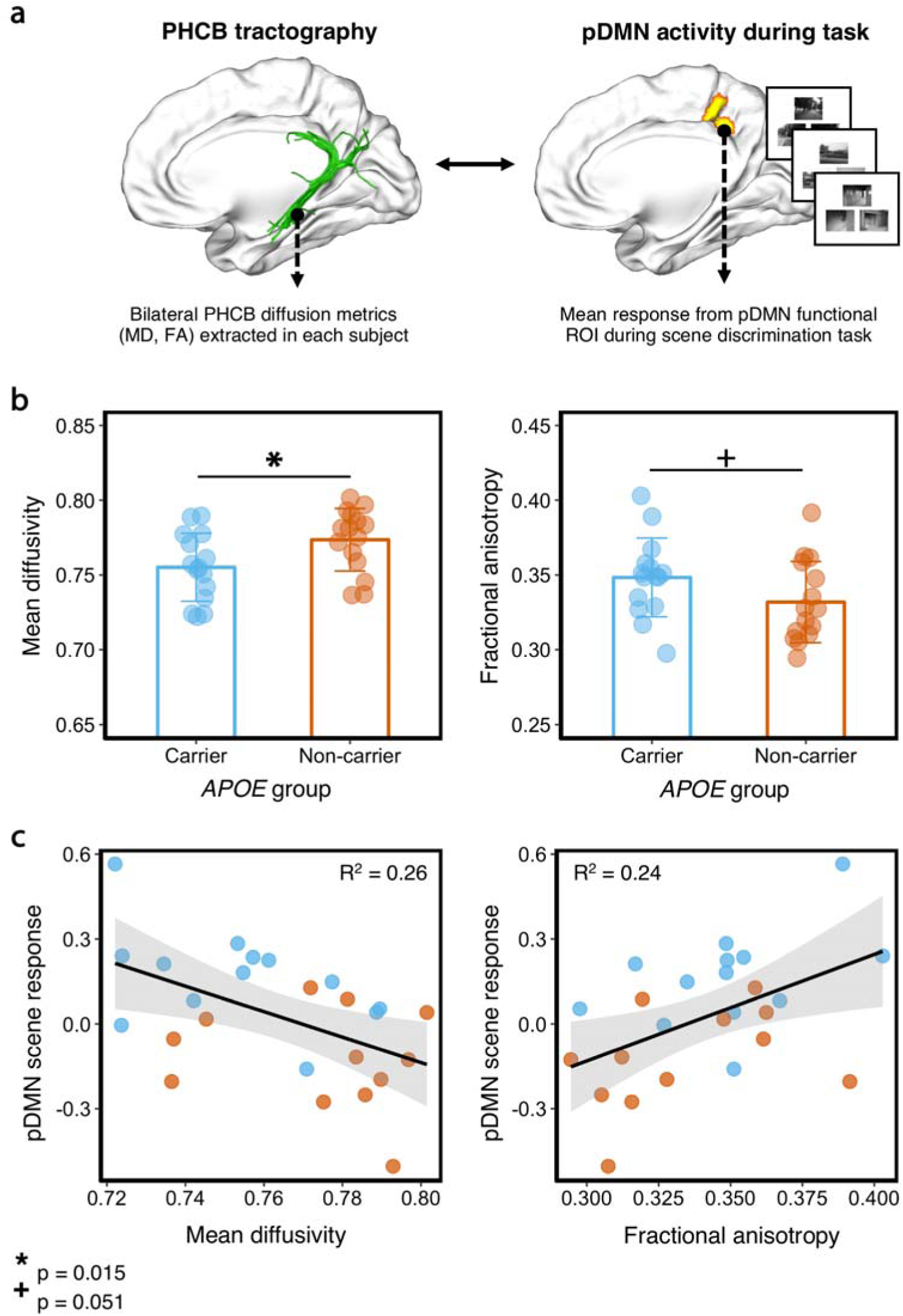
Comparing parahippocampal cingulum bundle (PCHB) tissue microstructure between *APOE*-ε4 carriers and non-carriers. (a) Left: Deterministic tractography was conducted in each subject and free water corrected indices of bilateral PHCB microstructure (MD, FA) were extracted (left). Right: To examine associations with functional activity, these metrics were correlated with BOLD activity from an independently-defined posterior default mode network (pDMN) functional region-of-interest during a perceptual discrimination task (Shine et al., 2015). Example scene trials for the perceptual ‘odd-one-out’ discrimination task are shown. (b) Plots comparing mean bilateral PHCB MD and FA for *APOE*-ε4 carriers and non-carriers. Individual data points are displayed jittered on each bar. (c) Scatter plots showing the association between scene (vs. “size” baseline) activity in pDMN and MD (left) and FA (right) in the PHCB. A total of 25 data points are shown on each scatter plot (13 carriers, blue markers; 12 non-carriers, orange markers; see Section 2.6).

#### 2.4.1. Parahippocampal cingulum reconstruction

Reconstruction of the PHCB followed a previously published and reliable protocol (termed “restricted parahippocampal cingulum”; see (Jones et al., 2013)). Following tract reconstruction in both hemispheres, the partial volume corrected maps for FA and MD were intersected with the PHCB tract masks to obtain mean bilateral measures of tract microstructure (MD, FA).

#### 2.4.2. Analysis of tractography data

MD and FA values of the bilateral PHCB in *APOE*-ε4 carriers and non-carriers were compared directly using directional Welch t-tests in R. We also report Default JZS Bayes Factors for our key analyses, computed using JASP (https://jasp-stats.org). The Bayes factor, expressed as BF_10_, reflects the strength of evidence that the data provide for the alternative hypothesis (H1) relative the null (H0). A BF_10_ much greater than 1 allows us to conclude that there is substantial evidence for the alternative versus the null hypothesis (Wagenmakers et al., 2017).

### 2.5 Tract-Based Spatial Statistics (TBSS)

Voxel-wise statistical analysis of the dMRI data was carried out using TBSS (Smith et al., 2006). This method involves non-linearly projecting subjects’ free water corrected statistical maps (both MD & FA) onto a mean tract skeleton and then applying voxel-wise cross-subject statistics. We applied a general linear model (GLM) contrasting *APOE*-ε4 carriers and non-carriers for each dMRI metric. To restrict our analysis to the PHCB, we extracted the PHCB mask from the Johns Hopkins University ICBM-DTI-81 white-matter atlas using FSLview (e.g., (Heise et al., 2014)). Significant clusters were extracted using Threshold-Free Cluster Enhancement (Smith and Nichols, 2009) with a corrected alpha of p = 0.05. Additional exploratory whole brain analyses were conducted using the same TFCE-corrected statistical threshold. All reported coordinates are in Montreal Neurological Institute (MNI-152) space.

### 2.6. Functional MRI methods

Further information regarding fMRI acquisition, preprocessing and analysis can be found in Shine et al. (Shine et al., 2015). Five participants were excluded from the fMRI analysis due to subject motion (4 subjects) and scanner error (1 subject) resulting in a final sample of 25 participants (13 carriers & 12 non-carriers). The fMRI measure-of-interest was the BOLD response (percent signal change) in the pDMN during a perceptual ‘odd-one-out’ discrimination task for scenes and faces (Figure 1A). The pDMN ROI was defined independently using a different cognitive task (short-term memory); this analysis confirmed a significant group difference (carriers > non-carriers) during scene short-term memory in pDMN (Shine et al., 2015). Individual percent signal change values for scenes and faces (each against a “size” oddity baseline condition) were calculated from this ROI using FSL and correlated with diffusion metrics using directional Pearson’s r correlations. Directional Bayes factors and 95% Bayesian credibility intervals (BCI) are reported for all correlations. BCIs inform us that, given our observed data, there is a 95% probability that the true value of our effect (Pearson’s r) lies within this interval. Correlations between each fMRI task condition and PHCB microstructure were compared using a one-tailed Steiger Z test of dependent correlations in ‘cocor’ (http://comparingcorrelations.org/) (Diedenhofen and Musch, 2015).

## 3. Results

### 3.1. Comparing PHCB microstructure using tractography

*APOE*-ε4 allele carriers had significantly lower MD compared to non-carriers (t (28) = 2.3 p = 0.015, d = 0.84, BF_10_ = 4.55; Figure 1B). While there was a strong trend for PHCB FA in the predicted direction, the between-group difference just failed to reach significance (t (28) = 1.69, p = 0.051, d = 0.62, BF_10_ = 1.83; Figure 1B).

Given suggested gender differences associated with *APOE*-ε4 (Heise et al., 2014; Ungar et al., 2014), we also conducted this analysis without male participants (removal of one individual from each group). A significant difference was found between carriers and non-carriers for PHCB MD, though with a slightly larger effect size (t (26) = 2.42, p = 0.012, d = 0.92, BF_10_ = 5.5). A significant difference was also found for PHCB FA (t (26) = 2, p = 0.03, d = 0.75, BF_10_ = 2.82).

### 3.2. Voxel-wise approach

TBSS analyses identified a significant cluster in right posterior PHCB for FA (p= 0.02; 29, −49, −1), reflecting higher FA in *APOE*-ε4 carriers (Figure 2) - consistent with the tractography analysis. We found no TFCE-corrected clusters for MD. Using an uncorrected threshold of p = 0.005 (Postans et al., 2014), we identified a significant cluster in left posterior PHCB reflecting lower MD in carriers (p < 0.001; −28, −58, 0). An exploratory whole brain analysis (TFCE-corrected) revealed no significant clusters for either metric.

**Figure 2.**
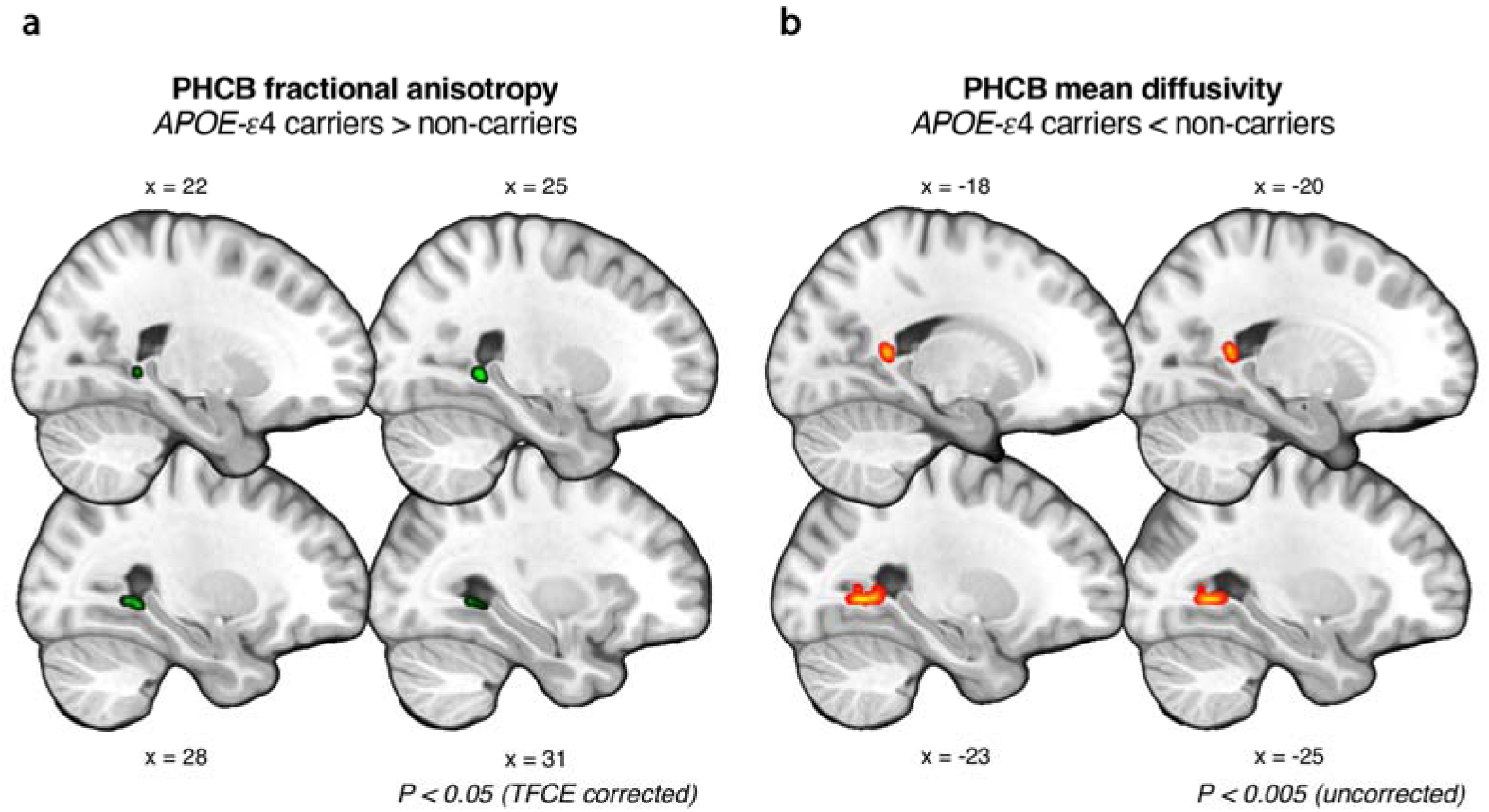
Comparing parahippocampal cingulum bundle (PHCB) microstructure in *APOE*-ε4 carriers and non-carriers using tract-based spatial statistics. (a) A significant cluster (shown in green) was found showing greater FA in *APOE*-ε4 carriers versus non-carriers in posterior PHCB (p < 0.05, TFCE-corrected). (b) A sub-threshold cluster (shown in red-yellow) reflecting lower MD in *APOE*-ε4 carriers versus non-carriers was identified in posterior PHCB (p < 0.005, uncorrected). For visualization purposes, clusters have been ‘thickened’ using ‘TBSS fill’ in FSL. There were no voxel-wise differences for MD that survived stringent correction.

### 3.3. The relationship between pDMN activity and PHCB microstructure

To examine the functional relevance of these structural connectivity metrics, we tested whether inter-individual variation in PHCB microstructure (MD, FA) was associated with fMRI response in the pDMN during an ‘odd-one-out’ discrimination task for scenes and faces (Shine et al., 2015). Across all subjects, we found a significant negative association between PHCB MD and scene activity (vs. “size” baseline) in the pDMN (r = −0.51, p = 0.01, BF_10_ = 12.1, 95% BCI [−0.73, −0.13]; Figure 1C). There was no significant association between MD and face activity (r = −0.03, p = 0.01, BF_10_ = 0.29, 95% BCI [−0.45, −0.01]). A one-tailed Steiger Z test revealed a significant difference between these coefficients (z = 2.5, p < 0.01). For PHCB FA, we likewise observed a significant association with scene, but not face, pDMN BOLD response (scene: r = 0.49, p = 0.01, BF_10_ = 8.87, 95% BCI [0.12, 0.72]); face: r = 0.12, p = 0.01, BF_10_ = 0.41, 95% BCI [−0.45, −0.01]; Figure 1C). The correlation between PHCB FA and scene activity was significantly greater than the correlation with face activity (z = 2, p = 0.02).

## 4. General discussion

Based on the view that pDMN vulnerability to amyloid-ß arises from its role as a large-scale connectivity hub (Bero et al., 2012; Brown et al., 2011; Buckner et al., 2009; de Haan et al., 2012; Jagust and Mormino, 2012), we asked whether young adults at heightened genetic risk for AD (via presence of the *APOE*-ε4 allele) would show increased pDMN structural connectivity (Greicius et al., 2009). Supporting this hypothesis, we found that *APOE*-ε4 carriers, relative to non-carriers, had microstructural differences in the PHCB – a white matter tract linking the pDMN with the MTL, particularly parahippocampal regions (Heilbronner and Haber, 2014). Moreover, inter-individual variation in PHCB microstructure was selectively associated with pDMN activity during a scene discrimination task that is sensitive to early AD (Lee, 2006).

The pDMN has been labelled the brain’s epicenter (Hagmann et al., 2008), given its disproportionately high structural/functional connectivity (Buckner et al., 2009; Hagmann et al., 2008). This region is also one of the first brain areas to undergo amyloid-ß deposition in AD (Gonneaud et al., 2016; Palmqvist et al., 2017). The early deposition of amyloid-ß in pDMN suggests that the high connectivity/activity demands on this region may, over the lifespan, lead to amyloid-ß accumulation and ultimately atrophy and cognitive decline (Bero et al., 2011). In human neuroimaging studies, strong within-subject correspondence has been found between pDMN functional connectivity strength and subsequent amyloid-ß load in individuals MCI (Myers et al., 2014). Elevated pDMN connectivity in low-amyloid individuals (Aß-) has also been associated with increased amyloid-ß deposition at follow-up (Jack et al., 2013). These increases in functional connectivity in older individuals, however, could reflect a compensatory response induced by early pathology (Jagust and Mormino, 2012; Jones et al., 2016; Schultz et al., 2017).

In young adult *APOE*-ε4 carriers, who are highly unlikely to harbor amyloid-ß, increased functional activity in pDMN and MTL has been seen across AD-relevant cognitive tasks (Dennis et al., 2010; Filippini et al., 2009; Shine et al., 2015). Young *APOE*-ε4 carriers also display greater intrinsic functional connectivity in the DMN compared to non-carriers (Filippini et al., 2009) - consistent with the view that functional activity differences may reflect increased connectivity. This contrasts with studies in older, cognitively-normal *APOE*-ε4 carriers, which report decreased functional connectivity (and also activity) in pDMN regions (Sheline et al., 2010).

Extending these studies, we found that college-aged *APOE*-ε4 carriers had increased *structural* connectivity (see below) in the PHCB – the main white matter pathway of the pDMN (Greicius et al., 2009). Specifically, young adult *APOE*-ε4 carriers had lower MD and higher FA compared to non-carriers. The direction of this effect contrasts with studies in older, cognitively-normal *APOE*-ε4 carriers and MCI, where decreased FA (and increased MD) is typically seen (Heise et al., 2014; Villain et al., 2008). Further, to demonstrate that these differences in structural connectivity are linked to difficulties modulating pDMN activity in *APOE*-ε4, we correlated inter-individual variation in PHCB microstructure with pDMN BOLD response during a scene discrimination task that is sensitive to early cognitive changes in AD [37]. This multi-modal, individual differences approach demonstrated that individuals with the highest pDMN activity during scene discrimination had the highest structural connectivity in the PHCB (lower MD/higher FA), suggesting that individual variation in structural connectivity in the PHCB may impact activity in pDMN, and subsequent vulnerability to amyloid-ß in later life (Jagust and Mormino, 2012). While group differences in MD and FA most likely reflect an impact of *APOE*-e4 on some aspect(s) of structural connectivity, we cannot readily determine what these are; variation in these diffusion metrics could arise from multiple, functionally-relevant biological properties (e.g., myelination, membrane permeability and/or axon number, diameter and voxel-wise configuration (D K Jones et al., 2013)).

One interpretation of these white matter differences is that they reflect early neuropathology, such as axonal loss or demyelination. Studies in asymptomatic adult carriers of a disease-causing *PSEN1* mutation (Ryan et al., 2013), and also in older adults with a parental history of AD (Melah et al., 2016), have reported, like here, greater FA (and lower MD) in a wide range of tracts, including the cingulum. Greater FA has also been observed in individuals with Aβ+ versus Aβ-MCI (Racine et al., 2014). Such differences have been attributed to axonal loss within specific fiber sub-populations of complex crossing-fiber areas (Douaud et al., 2011). This would lead to a reduction in fiber complexity and a concomitant increase in local anisotropy. Given that these *APOE*-ε4 carriers will not harbor amyloid-ß (Mormino, 2014), however, this neuropathological interpretation seems highly unlikely. Further, the pattern reported here is opposite to that seen typically in older individuals, where studies have shown reported lower FA/higher MD in individuals with AD and MCI ((Mito et al., 2018; Rieckmann et al., 2016) but see (Racine et al., 2014)). Rather, these findings support a LSV view of AD risk, where early-life, non-pathologically driven structural and functional alterations in specific brain networks may confer risk for later-life AD neuropathology (Jagust and Mormino, 2012).

Given this, one possible explanation for these white matter differences is that *APOE*-ε4 carriers and non-carriers undergo different patterns of white matter maturation. Previous studies suggest that efficient communication between distributed brain regions may emerge across development via overgrowth and then pruning of redundant axons (Yeatman et al., 2012). *APOE*-ε4 carriers, therefore, may show somewhat delayed axonal pruning of the late-maturing cingulum (Yeatman et al., 2014) during a critical period, such as adolescence (Yeatman et al., 2012), which leads to an overshoot in tissue microstructure and relative increases in pDMN neural activity (Figure 4). This increased pDMN activity in young adult *APOE*-ε4 carriers (as seen here during scene discrimination) may thus reflect some form of lifelong reduced network efficiency (Jagust and Mormino, 2012) or flexibility (Westlye et al., 2011), which impacts the ability of the pDMN to modulate activity (or state-dependent connectivity with MTL (Harrison et al., 2016; Westlye et al., 2011)) required to accommodate the need of a particular task.

Critically, these early life increases in pDMN structural connectivity (i.e., higher FA/lower MD), and concomitant changes in functional activity (Shine et al., 2015), may portend a faster decline in connectivity over the lifespan (Brown et al., 2011; Felsky, 2013), which ultimately leads to early amyloid-ß deposition and neurodegeneration (de Haan et al., 2012) (depicted in Figure 4). For instance, a cross-sectional study, which applied graph theory to measure the network characteristics of dMRI data, found that younger *APOE*-ε4 carriers had greater ‘local interconnectivity’ relative to non-carriers but exhibited a steeper age-related reduction ((Brown et al., 2011); see also (Felsky, 2013)). A potential compensatory increase in connectivity/activity, in response to accumulating amyloid-ß pathology (Schultz et al., 2017), will result in nodal stress and ultimately network failure (Jones et al., 2016), as reflected by a second steep decline in network integrity (activity/connectivity; Figure 3). A recent study, for instance, found that amyloid-ß contributes to the spreading of tau pathology via the PHCB (Jacobs et al., 2018). Future multi-modal imaging studies, conducted longitudinally across the lifespan, would provide further insights into how *APOE*-ε4 influences white matter microstructure and task-related activity across the lifespan.

**Figure 3.**
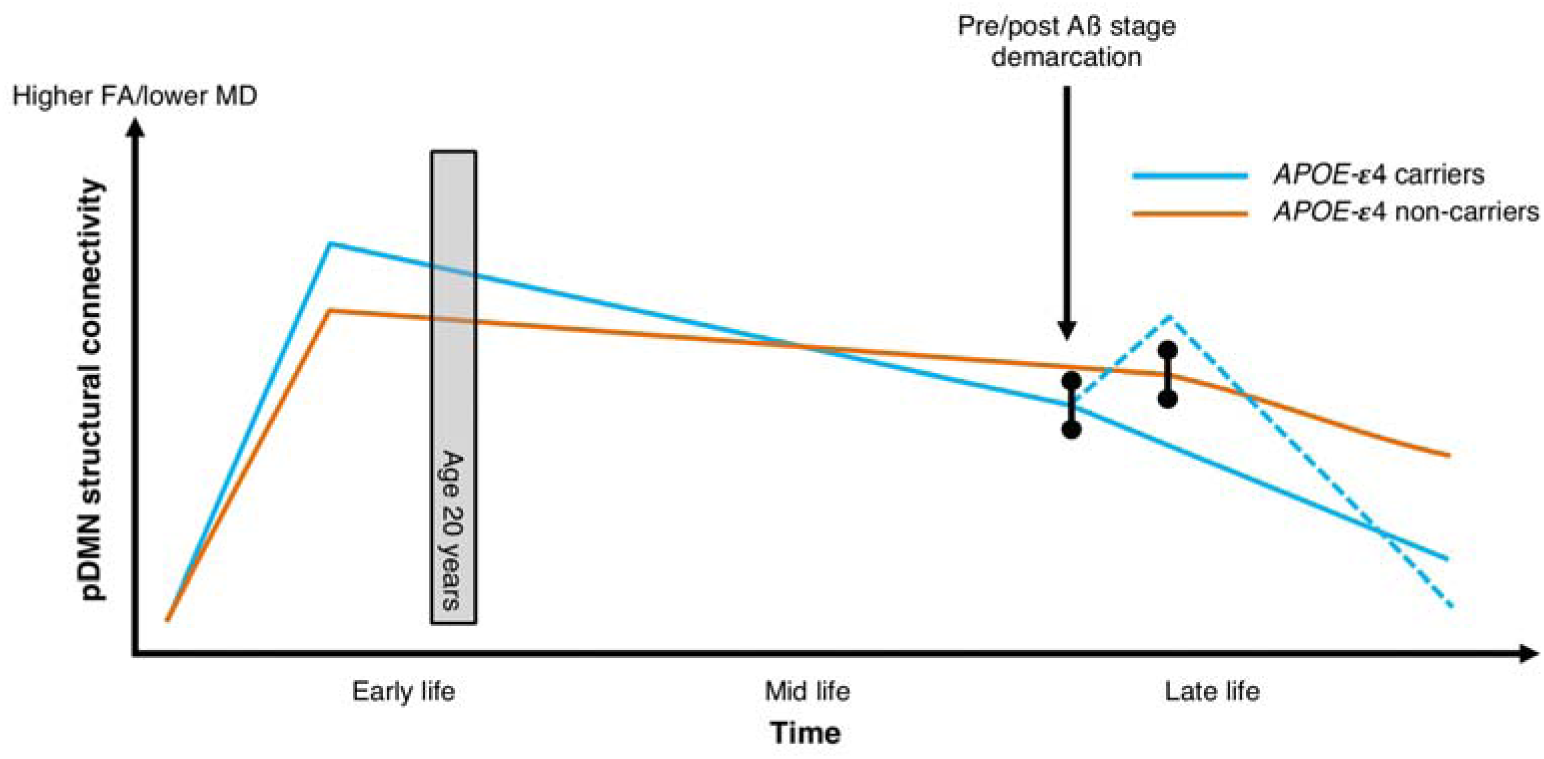
Posterior DMN structural connectivity and amyloid deposition across the lifetime. Figure *depicts* hypothetical trajectories of pDMN structural connectivity in *APOE*-ε4 carriers and non-carriers (as can be quantified using microstructural metrics derived from dMRI – i.e., FA and MD). Increased structural connectivity in young adult *APOE*-ε4 carriers (the age of our sample indicated by a grey box at 20 years) - which emerges over development – is proposed to lead to steeper decline across the lifespan (Brown et al., 2011; Felsky, 2013). Variation in structural connectivity across the lifespan leads to different demarcation points for amyloid-ß aggregation, with *APOE*-ε4 carriers showing earlier accumulation. The dashed blue indicates a hypothesized increase in connectivity in response to initial amyloid-ß burden – which may be mirrored in activity changes (Jagust and Mormino, 2012). Amyloid-ß deposition leads to “wear and tear” in the pDMN and a steep later-life decline in network structural connectivity, and eventual network failure (Jones et al., 2016).

While comparable to previous studies (Dennis et al., 2010; Filippini et al., 2009; Oh and Jagust, 2013), the sample size in the current study is relatively modest. This issue is partly mitigated by a clear hypothesis-driven approach (Button et al., 2013) and Bayesian analyses showing that our findings have high evidential value (Dienes, 2014).

While we observed significant differences for both FA and MD, our reported effects were somewhat stronger for MD, particularly for the tractography analysis. This is consistent with reports that FA shows greater intra-tract variability than MD – i.e. tracts do not have a signature FA value that is consistent along the tract length (Yeatman et al., 2012). Future dMRI studies using advanced tract profiling and biophysical modelling would shed further insight into the relation between *APOE*-e4 and PHCB microstructure (Assaf et al., 2017; Yeatman et al., 2012).

## Conclusion

To conclude, we have shown that *APOE*-ε4-related increases in pDMN activity (Shine et al., 2015) are linked to structural connectivity in the PHCB - the main white matter conduit linking pDMN with the MTL (Heilbronner and Haber, 2014). Specifically, *APOE*-ε4 carriers had significantly lower MD, and higher FA, in this pathway – the opposite effect to that seen in cognitively-normal older *APOE*-ε4 carriers (Felsky, 2013). By combining dMRI and BOLD fMRI measures, we showed that inter-individual variation in PHCB microstructure (increased FA/decreased MD) was linked to increased pDMN activity during a scene discrimination task that is affected in AD (Lee, 2006). These findings support a LSV model of AD risk, whereby connectivity-associated increases in pDMN activity across the lifespan may confer risk for amyloid-ß accumulation in later life – one of the earliest biomarkers of AD pathology.

## Author contributions

CJH, ADL and KSG designed research; CJH collected the data; CJH, JPS, HW and MP analyzed the neuroimaging data; RS and JW analyzed the genetic data; CJH wrote the paper with support from all other authors; CJH and JPS are joint first authors.

## Acknowledgments

This work was supported by Alzheimer’s Research UK (KSG), the Medical Research Council (MR/N01233X/1: KSG; G1002149: KSG, CJH), a Wellcome Trust Strategic Award (104943/Z/14/Z; CJH), The Waterloo Foundation, the Biotechnology and Biological Sciences Research Council (BB/I007091/1; KSG, MP), and the Welsh Government (via the Wales Institute of Cognitive Neuroscience). We would like to thank John Evans, Martin Stuart and Peter Hobden for scanning support.

